# Pain to remember: a single incidental association with pain leads to increased memory for neutral items one year later

**DOI:** 10.1101/035212

**Authors:** G. Elliott Wimmer, Christian Büchel

## Abstract

Negative and positive experiences can exert a strong influence on later memory. Our emotional experiences are composed of many different elements – people, place, things - most of them neutral. Do emotional experiences lead to enhanced long-term for these neutral elements as well? Demonstrating a lasting effect of emotion on memory is particularly important if memory for emotional events is to adaptively guide behavior days, weeks, or years later. We thus tested whether aversive experiences modulate very long-term episodic memory in an fMRI experiment. Participants experienced episodes of high or low pain in conjunction with the presentation of incidental, trial-unique neutral object pictures. In a scanned surprise immediate memory test, we found no effect of pain on recognition strength. Critically, in a follow-up memory test one year later we found that pain significantly enhanced memory. Neurally, we provide a novel demonstration of activity predicting memory one year later, whereby greater insula activity and more unique distributed patterns of insular activity in the initial session correlated with memory for pain-associated objects. Generally, our results suggest that pairing episodes with arousing negative stimuli may lead to very long-lasting memory enhancements.

## Introduction

Episodic experiences that lead to negative consequences, resulting in pain, fear, anger, and other aversive emotions, may remain in our memories longer than non-emotional experiences. In some cases, memories for negative events can even impair everyday life, as in post-traumatic stress disorder (Shin and Liberzon, 2010). A negative episode contains many separate elements that are often by themselves neutral. Prioritizing the content of emotionally arousing experiences in memory may reflect an adaptive function (Ochsner, 2000). When later encountering neutral stimuli that were present during an emotional episode, attentional orienting may be more rapid, allowing detection of new potential threats; at the same time, recognition may facilitate the recall of further information about relevant past experiences. Importantly, however, research to date on the emotional modulation of neutral stimuli has not revealed a consistent enhancement of memory (Phelps et al., 1997; Maratos and Rugg, 2001; Smith et al., 2004a).

A rich literature has studied the modulation of memory by emotion, using stimuli such as well-characterized affective pictures (for review, see Reisberg and Heuer, 2004; LaBar and Cabeza, 2006). In several studies, a benefit for remembering emotional stimuli has been found weeks or even a year later (Bradley et al., 1992; Cahill et al., 1996; Ochsner, 2000; Dolcos et al., 2005; Wagner et al., 2006; Weymar et al., 2011). However, the use of emotional stimuli in memory studies presents several difficulties. Generally, emotional stimuli can trigger an approach or avoidance response without the need for memory of any previous experience; in contrast, neutral elements of emotional experiences cannot elicit the same automatic response without memory. Also, emotional stimuli have been shown to attract increased attention during initial encoding that, along with increased semantic relatedness, may account for observed memory benefits (Talmi and McGarry, 2012; Talmi, 2013). Further, in a memory test, reexposing participants to emotional stimuli can lead to new emotional processing, making neural effects difficult to interpret. At the same time, a memory test may allow for compounding effects of attention that could further influence later memory.

To avoid these concerns, researchers have utilized designs where neutral items are associated with emotional or non-emotional contexts during encoding and then memory is tested for the neutral items (Phelps et al., 1997; Maratos and Rugg, 2001; Smith et al., 2004a). When encoding is incidental (as in the majority of everyday experience), studies have reported null effects or even emotional memory impairments, except when participants are required to generate a semantic connection between the emotional context and the neutral item (Maratos and Rugg, 2001; Erk et al., 2003; Smith et al., 2004a; Smith et al., 2004b; Smith et al., 2006; Bingel et al., 2007; Forkmann et al., 2013; Zhang et al., 2015). Importantly, consolidation processes are known to play an important role in enhancing memory for emotional experiences, allowing for memory strengthening via neuromodulatory-induced plasticity (McGaugh, 2004; Yonelinas and Ritchey, 2015). Currently, however, only one study of several has reported increased day-later incidental memory for neutral stimuli associated with negative emotions (Jaeger et al., 2009; Jaeger and Rugg, 2012; Schwarze et al., 2012).

Consolidation intervals longer than one day may reveal effects on emotion and memory that are not apparent in immediate or next-day memory, but no previous studies have examined the effects of extended consolidation on incidental memory for neutral stimuli. Demonstrating a lasting effect of emotion on memory is of particular interest if memory for emotional events is to adaptively guide behavior days, weeks, or years later. Further, previous studies on very long-term emotional memory have not shown whether neural activity during the initial experimental session predicts very longterm emotional memory (Dolcos et al., 2005). We thus investigated whether very longterm memory for incidental neutral stimuli is modulated by a single aversive association (Fig. 1a). In the initial fMRI session, neutral objects were presented once, incidentally paired with high or low pain (Fig. 1b). A scanned surprise memory test followed. One year later, participants returned to the lab for a follow-up memory test, allowing us to examine whether memory for the neutral objects was modulated by a single aversive experience one year before. Memory for affective experiences such as negative emotional pictures has been related to activity in the medial temporal lobe (MTL) including the amygdala (Murty et al., 2010). However, for pain, previous neuroimaging studies have found an overlap in insula activity during pain and short-term remembered pain (Albanese et al., 2007; Fairhurst et al., 2012). Further, studies of in post-traumatic stress disorder suggest a role for the insula in representing traumatic memories (Liberzon and Martis, 2006). Based on these findings and a hypothesized role of the anterior insula in processing the emotional and evaluative aspects of pain (Kurth et al., 2010; Wiech et al., 2014), we predicted that anterior insula activity during the initial experimental session may be related to very long term memory.

**Fig. 1.**
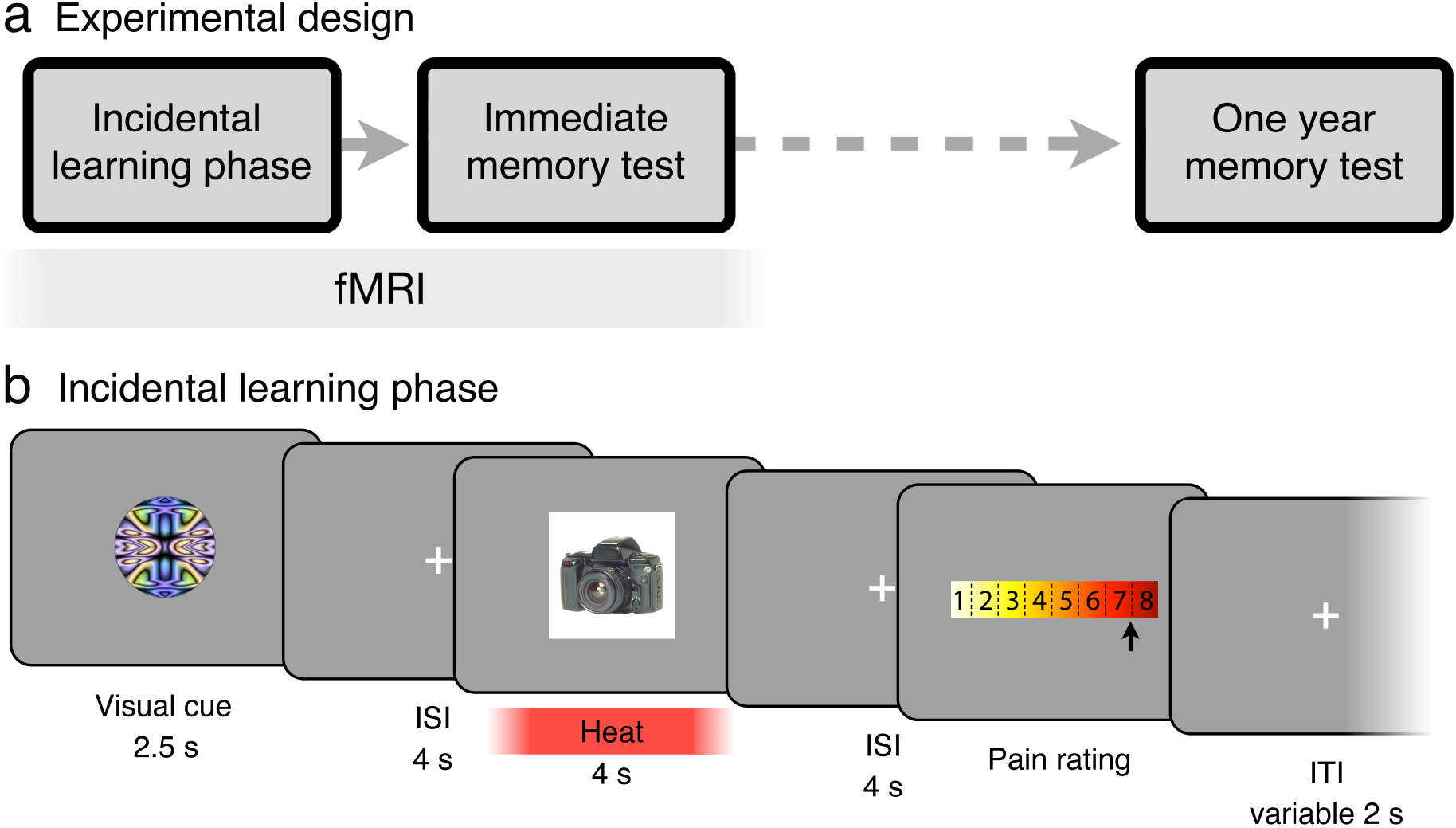
Pain and incidental long-term memory experiment. (a) Experimental design: The incidental learning phase was followed by a surprise memory test phase during fMRI scanning, in which participants responded with whether objects had been associated with high or low pain and then rated their recognition strength (Fig. S1a). One year later, participants returned to the lab for a follow-up surprise recognition memory test (Fig. S1b). (b) In the incidental learning phase, participants experienced high or low heat pain while being exposed to an incidental trial-unique object pictures. Participants then rated their experienced level of pain.

## Materials and Methods

**Participants:** A total of 31 subjects participated in the experiment. Participants were right-handed fluent German speakers with no self-reported neurological or psychiatric disorders and normal or corrected-to-normal vision. Data from 2 participants were excluded due to technical problems with the thermode and 5 participants were unable to return for the one-year follow-up behavioral test, leaving 24 participants (12 female; mean age, 25.8 years; range, 20-33 years). In one participant, pain memory confidence ratings and memory recognition strength in the immediate test session were not recorded due to a technical error; this participant was excluded from analyses using immediate session data. The one-year later follow-up session was conducted on average 362.9 days after the initial scanning session (range: 316-469 days). The Ethics committee of the Medical Chamber Hamburg approved the study and all participants gave written consent.

Analyses and results focus on the modulation of recognition memory strength by pain in the immediate and one-year tests. Results related to memory for pain (high vs. low) will be reported separately.

**Heat calibration**. Before the incidental learning phase, heat levels were calibrated for each subject to achieve the same subjective high and low aversive pain experience across subjects. Thermal stimulation was delivered via an MRI compatible 3 × 3 cm Peltier thermode (MSA; Somedic, Sweden), applied to the inner left forearm. During the visual presentation of a white square, heat was applied for 10 s. For pain rating, we used a 1-8 rating scale with 0.5-point increments, superimposed on a yellow-to-red gradient. An arrow cursor was moved from the initial mid-point starting location using left and right key-presses and ratings were confirmed with the space bar. A rating of ‘8’ corresponded to the highest level of heat pain a subject could endure multiple times. If the level of pain was intolerable, subjects moved the rating past the ‘8’ end of the scale, at which point a ‘9’ appeared on the screen. Subjects rated the pain associated with a pseudo-random list of 10 different temperatures ranging from 39.5 to 49.5°C. A linear interpolation algorithm then selected a low temperature estimated to yield a ‘2’ rating and a high temperature estimated to yield a ‘7.5’ rating.

**Procedure: incidental learning phase**. In the incidental learning phase, participants experienced high or low heat pain while being exposed to trial-unique incidental object pictures (Fig. 1b). Importantly, the encoding of the object pictures was incidental (not instructed), to more closely match the incidental nature of encoding in many real-world situations. Across 4 blocks, 33 high heat trials and 35 low heat trials were presented (Fig. 1b). On each trial a visual cue was presented for 2.5 s signaling likely high or low heat. After a 4 s ISI, the incidental object appeared. To allow for a better match between the appearance of the object and the onset of noticeable heat, heat onset started 0.75 s prior to object appearance (for a similar method, see Forkmann et al., 2013). The incidental object was presented for a total duration of 10 s, after which the temperature returned to baseline (33°C) over several seconds. After a 4 s ISI, the pain rating scale appeared. Subjects used the left and right buttons to move a selection arrow from the initial cursor position (randomized between 4.5-5.5) to their experienced pain level and pressed the down button twice to make their selection; responses were self-paced. After the subject entered their response, trials were followed by a variable 2 s mean (range: 0.5-6 s) inter-trial-interval (ITI).

To maintain attention on the screen during visual cue presentation, on a random 50% of trials the visual cue illumination flickered (decreased in illumination) once for 0.35 s. Flicker timing was randomly distributed throughout the first 1.5 s of visual cue presentation. Similarly, on a separately determined random 50% of trials the object picture flickered in illumination during heat stimulation. When either a visual cue or object flicker was detected, subjects were instructed to press the down button.

Two pseudo-random orderings of incidental object pictures were used for counterbalancing object and heat associations. The assignment of abstract circles to high and low heat was also counterbalanced across subjects, and after the first two blocks of the experiment, two new abstract circles were used as cues, with visual and verbal instruction about the new cues preceding the block. To investigate effects of anticipation and expectation violation, visual cues were probabilistically associated with the level of heat, correctly predicting high or low heat on 67% of trials (Atlas et al., 2010). On invalid trials, the alternative heat level was administered. Additionally, 6 trials included a probe of cue-related pain expectancy: after 2.5 s of cue presentation, a question appeared below the cue asking subjects whether they expected low or high heat to follow. These probes were used to test the learning of the visual cue-pain associations. After the probe, trials continued as normal. During the three breaks between the four incidental learning phase blocks, the thermode was moved to a new location on the inner arm to avoid sensitization.

To maintain similar differences in subjective experience between the high and low heat conditions, temperatures were automatically adjusted throughout the task to maintain the targeted pain rating values. If the median of the previous 6 validly cued low heat trials fell below a rating of 1.5, the low temperature was increased by 0.2°C; if the median rating was above 3, the low temperature was decreased by 0.2°C. For the high temperature, if the median rating fell below 7.5, the high temperature was increased by 0.2°C (if the temperature was below 50.5°C). If a rating of “9” was given, indicating an intolerably high level of pain, the high temperature was decreased by 0.8°C.

**Procedure: immediate memory test phase**. In the scanned surprise memory test following the incidental learning session, we assessed memory for the level of pain administered with the object and recognition strength (Fig. S1a). (Results related to pain value memory will be reported separately.) Participants saw each of the 68 “old” objects from the incidental learning phase intermixed with 20 “new” objects (Fig. S1a). On each trial a single object was presented alone for 5 s. Next, after a 1 s ISI, an unmarked heat scale with superimposed left- and right-pointing arrows was shown. Subjects pressed the left or right buttons to indicate whether they thought that the object had been associated with low heat pain or high heat pain in the incidental learning phase. For objects that subjects definitely considered to be “new”, subjects were told that they could pick either the high or low heat response at random. If they were not sure an object was new, subjects were instructed to try to recall the level of heat it may have be associated with. All test phase responses were self-paced. Next, a confidence rating screen appeared with 4 levels of response: “guess”, “somewhat certain”, “certain”, and “very certain”. For stimuli subjects believed were definitely new, subjects were instructed to respond with a low confidence answer. After a variable ISI (mean: 4 s; range: 3-6.5 s), the 6-point memory recognition strength scale was presented. Subjects indicated whether they thought the object was “new” (not previously seen) or “old” (seen during the learning task) with 6 levels of response: “certain new”, “somewhat certain new”, “guess new”, “guess old”, “somewhat certain old”, “certain old”. Subjects used the left and right buttons to move from the randomly initially highlighted “guess new” or “guess old” response option to their selected response and then pressed the down button twice to make their selection. A variable ITI with a mean of 4 s (range: 2-8 s) followed. The order of the old pictures was pseudo-randomized from the incidental learning phase order, and the old and new pictures were pseudo-randomly intermixed.

**One year later memory test phase**. Approximately one year after the initial fMRI experimental session, participants returned to the lab to complete a surprise behavioral memory test session (Fig. S1b). On each trial, objects were displayed alone for 3 s. Then, participants rated their recognition strength for the object on the 1-6 new-to-old scale. After a 1 s ISI, for objects rated as “old” participants then indicated whether they thought the object had been incidentally paired with pain in the incidental learning session. For objects rated “new” participants waited for a 6 s ISI. A variable 3 s mean ITI followed each trial. Participants saw each of the 68 old objects from the incidental learning phase one year prior intermixed with 32 new objects that had not been seen in the experiment before.

The duration and distribution of ITIs (or “null events”) was optimized for estimation of rapid event-related fMRI responses as calculated using Optseq software (http://surfer.nmr.mgh.harvard.edu/optseq/). The experimental tasks were presented using Matlab (Mathworks, Natick, MA) and the Psychophysics Toolbox (Brainard, 1997). The year-later behavioral session was completed on a laptop computer. The task was projected onto a mirror above the subject’s eyes. Responses were made using a 4- button interface with a “diamond” arrangement of buttons. At the end of the experiment, subjects completed a paper questionnaire querying their knowledge of the task instructions and their expectations (if any) regarding the incidental object pictures. Task instructions and on-screen text were presented in German; on-screen text was translated into English for the methods description and task figures.

**Behavioral Analysis**. The primary behavioral question was whether memory one year later was modulated by pain experience in the incidental learning session. Multilevel regression models as implemented in R (R-project.org) were used to investigate immediate and year-later recognition memory strength.

**fMRI Data Acquisition**. Whole-brain imaging was conducted on a Siemens Trio 3 Tesla system equipped with a 32-channel head coil (Siemens, Erlangen, Germany). Functional images were collected using a gradient echo T2*-weighted echoplanar (EPI) sequence with blood oxygenation level-dependent (BOLD) contrast (TR = 2460 ms, TE = 26 ms, flip angle = 80, 2 x 2 x 2 mm voxel size; 40 axial slices with a 1 mm gap). Slices were tilted approximately 30° relative to the AC–PC line to improve signal-to-noise ratio in the orbitofrontal cortex (Deichmann et al., 2003). Head padding was used to minimize head motion; no subject’s motion exceeded 3 mm in any direction from one volume acquisition to the next. For each functional scanning run, four discarded volumes were collected prior to the first trial to allow for magnetic field equilibration.

During the incidental learning phase, four functional runs of an average of 190 TRs (7 min and 48 s) were collected, each including 17 trials. During the memory test phase, four functional runs of an average of 196 TRs (8 min and 2 s) were collected, each including 22 trials. If a structural scan had not been collected for the subject at the center within the past 6 months, structural images were collected using a highresolution T1-weighted magnetization prepared rapid acquisition gradient echo (MPRAGE) pulse sequence (1 x 1 x 1 mm voxel size) between the incidental learning phase and the value memory test phase.

**fMRI analyses**. Preprocessing and data analysis was performed using Statistical Parametric Mapping software (SPM8; Wellcome Department of Imaging Neuroscience, Institute of Neurology, London, UK). Before preprocessing, individual slices with artifacts were replaced with the mean of the two surrounding timepoints using a script adapted from the ArtRepair toolbox (Mazaika et al., 2009). Images were slice-timing corrected, realigned to correct for subject motion, and then spatially normalized to the Montreal Neurological Institute (MNI) coordinate space by estimating a warping to template space from each subject’s anatomical image and applying the resulting transformation to the EPIs. Images were filtered with a 128 s high-pass filter and resampled to 2 mm cubic voxels. Images were then smoothed with a 6 mm FWHM Gaussian kernel.

fMRI model regressors were convolved with the canonical hemodynamic response function and entered into a general linear model (GLM) of each subject’s fMRI data. The six scan-to-scan motion parameters produced during realignment were included as additional regressors in the GLM to account for residual effects of subject movement.

fMRI analyses focused on whether activity during the incidental learning phase or memory test phase was correlated with immediate and one-year later measures of recognition memory strength. We first conducted “localizer” univariate analyses to identify main effects of pain in the incidental learning phase. The GLM included regressors for the cue (2.5 s duration), object and pain presentation (10 s duration), and the pain rating (variable duration). The cue regressor was accompanied by a modulatory regressor for high vs. low expected pain and the pain regressor was accompanied by a modulatory regressor for the pain rating given on that trial.

To examine the primary question about neural correlates of immediate and year-later recognition memory strength, we first examined activity during the incidental learning phase. We constructed two general linear models (GLMs): the first model focused on correlates of memory during the initial object presentation period, while the second model focused on correlates of memory during the peak pain period of the trial. The onset GLM included regressors for the cue period (0 s), pain onset (0 s), and pain rating onset (0 s). The cue period and the pain period regressors were accompanied by 4 parametric regressors entered in this order: immediate recognition memory strength for high pain objects, immediate recognition memory strength for low pain objects, one-year recognition memory strength for high pain objects, and one-year recognition memory strength for low pain objects. The peak pain GLM examined memory correlates once pain-related activation had reached a peak, as estimated using GLMs that systematically varied the onset of the pain rating regressor from in 1 s increments from 0 s to 8 s post-onset. We found peak responses in the anterior insula at 5 s post-onset, and thus we focused on this period. The peak pain GLM included regressors for the cue period (2.5 s), initial pain period (5 s), late pain period (5 s) and pain rating (2 s). The cue period and the pain period regressors were accompanied by the 4 parametric immediate and year-later memory regressors described above.

We next examined neural correlates of immediate and year-later memory during the surprise memory test phase. In the memory test phase, object pictures were presented alone for 5 s at the start of the trial and again during the memory recognition strength response. As the full trial concerned memory questions for the same object, and as cognitive and memory processes likely engaged some maintenance of the object even when the stimulus was not on the screen (between the initial presentation and the memory recognition rating), we modeled memory during the full trial duration. Trial durations varied based on individual response times. The test phase model thus included a regressor for the full trial, with 5 parametric regressors entered in this order: a control contrast of old vs. new objects, immediate recognition memory strength for high-pain associated objects, immediate recognition memory strength for low-pain associated objects, one-year recognition memory strength for high-pain associated objects, and one-year recognition memory strength for low-pain associated objects. The use of separate parametric modulators for high- and low-pain objects allows for second-level memory contrasts while controlling for main effects of pain. For univariate analyses, linear contrasts of univariate SPMs were taken to a group-level (randomeffects) analysis.

Finally, we used representational similarity analysis (RSA) to examine patterns of activity evoked by stimuli that were remembered vs. forgotten one year later during the initial fMRI session (Kriegeskorte et al., 2008). For these analyses, we modeled the non-smoothed fMRI data in GLMs with separate regressors for each trial, in addition to motion nuisance regressors as described above. In the incidental learning phase, the individual trial regressor duration covered the full 10 s of pain and object presentation. In the test phase, similar to the univariate GLM described above, the individual trial regressor duration covered the full memory trial, including the initial presentation period and the recognition strength response. These GLMs provided beta values for each voxel for each trial, which we extracted within regions of interest. Correlations between patterns of beta values in ROIs within pain level (e.g. high-pain remembered objects, high-pain forgotten objects) were computed using Pearson’s r. Resulting r values were Fisher-transformed to z-scores before statistical comparison.

Our primary RSA analysis focused on differences in representational similarity for high pain-associated objects related to memory one year later. Given the univariate memory-related effect in the test phase, our RSA analyses focused on test phase activity. Previous studies have shown that higher within-item similarity across repetitions or across encoding to retrieval are related to better memory (Xue et al., 2010; Ritchey et al., 2013). Given the relationship between within-item similarity and memory, we expected that successful memory may be related to more distinct processing of items, leading to higher across-item dissimilarity. A parallel control analysis was conducted using immediate recognition strength; as there were few instances of forgotten objects in the immediate test (old objects rated as new), recognition values were instead binned based on above- and below-mean immediate recognition memory strength. We also examined representational similarity across learning and test-phase presentations (Ritchey et al., 2013). However, initial control analyses indicated that within-item similarity was strongly affected by the application of pain: learning-test correlations in an object-responsive region of the visual cortex were highly significant when the initial learning phase was modeled with a 0 s duration regressor (and the initial test phase presentation was modeled with a 5 s duration regressor), but these within-item correlations were eliminated when the full pain period was modeled. Given the absence of within-item similarity effects, we did not conduct further memory-related analyses across learning to test.

**Regions of interest**. We report results corrected for family-wise error (FWE) due to multiple comparisons (Friston et al., 1993). We conduct this correction at the peak level within small volume ROIs for which we had an a priori hypothesis (after an initial thresholding of p < 0.005 uncorrected) or at the whole-brain cluster level, with a cluster threshold of 10 voxels. With the exception of pain-related activation (Table S1), we found no significant results outside of our a priori regions of interest.

We focused on a priori ROIs in the anterior insula and medial temporal lobe (MTL). For the anterior insula, we first created a bilateral anterior insula mask (Brooks et al., 2002; Wiech et al., 2014) covering the insular cortex anterior to y = 9 as well as several millimeters lateral of the insular cortex to account for signal blurring and anatomical variability. This mask was further restricted by the main effect of pain taken from the localizer GLM defined above, thresholded at p < 0.0001 uncorrected. The MTL ROI, including the hippocampus, parahippocampal cortex, and amygdala, was based on the AAL atlas (Tzourio-Mazoyer et al., 2002). All voxel locations are reported in MNI coordinates, and results are displayed overlaid on the average of all subjects’ normalized high-resolution structural images.

## Results

**Behavioral**. In the immediate surprise memory test phase, on each trial, participants first viewed an object picture for 5 s. Participants then indicated if the object had been associated with high or low heat pain and then rated their confidence. Next, participants rated their recognition strength on a 1-to-6 new-to-old scale (Fig. S1a). We found that participants reliably discriminated objects seen during incidental learning from new objects (t(_22_) = 16.60, p < 0.001; immediate memory rating data missing for one participant). However, pain did not affect immediate recognition memory (high heat objects: 4.94 ± 0.10 (mean ± SEM); low heat objects: 4.96 ± 0.10; new objects: 2.14 ± 0.13; t(_22_) = 0.41, p = 0.68; Fig. 2a). The immediate memory measure was also not related to pain ratings or administered temperature (regression analysis; pain ratings: p = 0.88; heat temperature: p = 0.82). These results support the previously reported absence of a memory enhancement for emotion-associated neutral stimuli when tested immediately (e.g. Maratos et al., 2001; Schwarze et al., 2012). The null effect of pain on immediate memory, as well as previous reports supporting an interruptive effect of pain on memory and cognitive processing (Bingel et al., 2007; Talmi and McGarry, 2012), suggest that pain did not increase attention to incidental objects in the incidental learning phase.

**Fig. 2.**
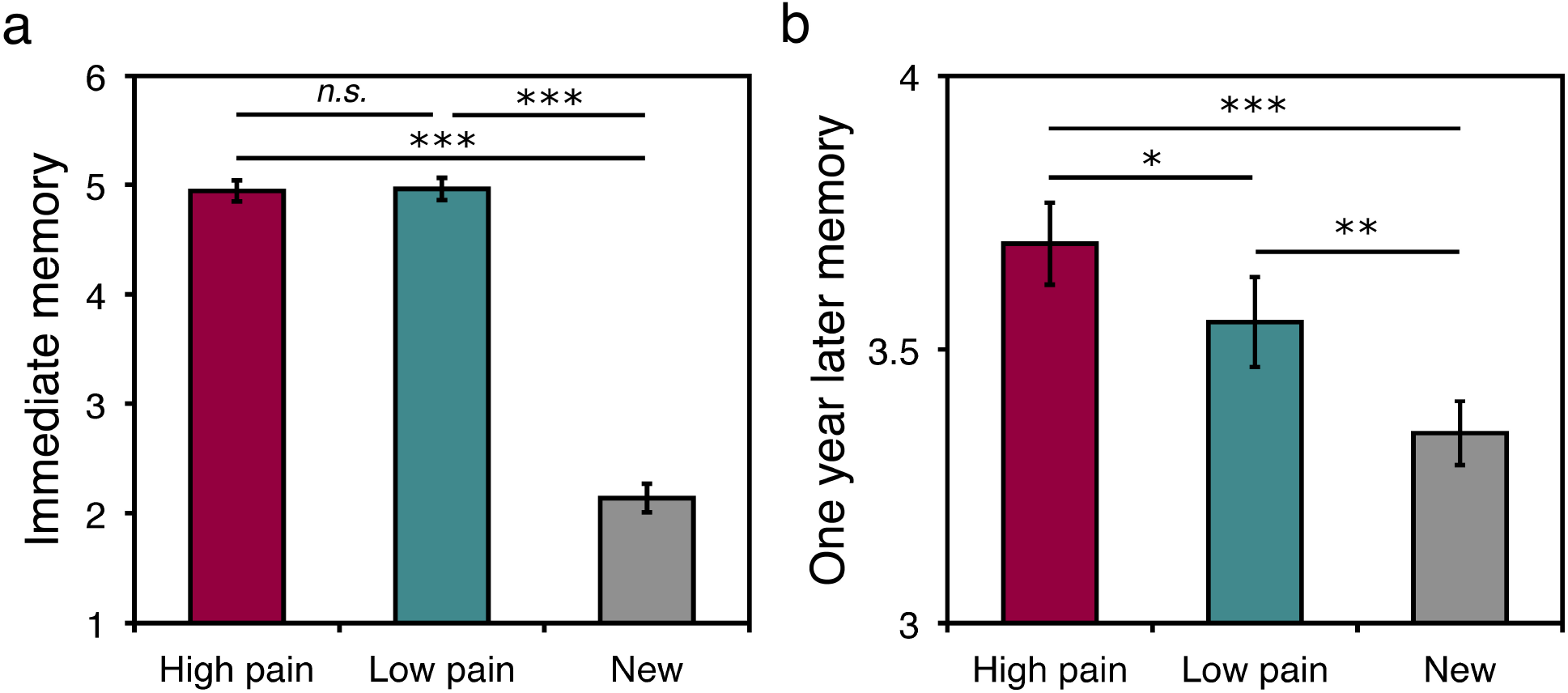
A single association with pain enhances memory for objects one year later. (a) Memory strength (rated on a 1-6 new-to-old scale) in the immediate memory test for high pain-associated objects, low pain-associated objects, and new objects revealed no difference due to pain experience. (b) After one year, pain-associated objects showed significantly higher memory than low pain-associated objects. Significance is indicated with asterisks: * p < 0.05; ** p < 0.01; *** p < 0.001.

One year after the initial fMRI session, participants returned to the lab for a behavioral surprise memory test session (Fig. S1b). This allowed us to answer the critical question of whether a single incidental pairing with pain affected memory for neutral items one year later. We indeed found that memory for high-pain associated objects was significantly higher than memory for low-pain associated objects and new objects (high heat objects; 3.69 ± 0.07; low heat objects: 3.55 ± 0.08; new objects: 3.34 ± 0.06; high vs. low, t(23) = 2.44, p = 0.023; high vs. new, t(23) = 6.67, p < 0.001; Fig. 2b). Memory for low-pain objects was also significantly higher than memory for new objects (t_(23)_ = 2.91, p = 0.0078; Fig. 2b). Controlling for initial memory ratings in the immediate test as well as initial pain value memory responses, we found that high vs. low pain remained a significant predictor of year-later memory (coef. 0.13 ± 0.06; t(20) = 2.21, p = 0.027). Importantly, on a trial-by-trial basis, initial recognition strength was unrelated to year-later memory (coef. 0.01 ± 0.02; t(_20_) = 0.64, p = 0.52). Pain value memory (“high pain” vs. “low pain” responses) showed a trending positive effect on year-later memory (coef. 0.11 ± 0.06; t(20) = 1.73, p = 0.08).

Additionally, pain ratings for the trial-by-trial pain experienced during individual objects positively predicted year-later memory (coef. 0.03 ± 0.01; t(_20_) = 2.32, p = 0.020; regression control for immediate session responses). The administered temperature of heat stimulation was also a significant predictor of year-later memory (coef. 0.02 ± 0.01; t(_20_) = 2.94, p = 0.0033). Notably, even within high-pain associated objects, temperature remained a significant positive predictor of year-later memory (coef. 0.02 ± 0.01; t(_20_) = 2.04, p = 0.042). The effect of heat temperature on memory suggests that the nociceptive and arousal-related responses due to variations in temperature are a robust predictor of the strength of consolidated memory after one year.

## fMRI

Does neural activity during the initial fMRI session correlate with year-later memory for pain-associated objects? In the incidental learning phase, we found no significant activation related to immediate memory for high-pain vs. low-pain associated objects (see Supplementary Results for uncorrected results). For memory one year later, during the incidental learning phase we also found no relationship between activity and memory for high-pain vs. low-pain associated objects, either at the onset of objects or during the peak pain period.

In the immediate memory test session, objects were presented in the absence of pain. We found no correlates of immediate recognition memory strength specific to high-pain associated objects or memory overall. Critically, we found a correlation between one-year later recognition memory strength for high-pain vs. low-pain associated objects and activity in the right anterior insula (Right, Anterior, Superior: 34, 24, 4; Z = 3.84, p = 0.036 SVC; Fig. 3). The peak of the insula cluster is within a region of insula activation correlated with trial-by-trial pain ratings in the incidental learning session (peak: 32, 14, 8; Z = 6.36, p < 0.001 whole-brain FWE; Table S1). In contrast, we found no significant correlates of memory across high- and low-pain associated objects or correlates of memory for low-pain objects alone.

**Fig. 3.**
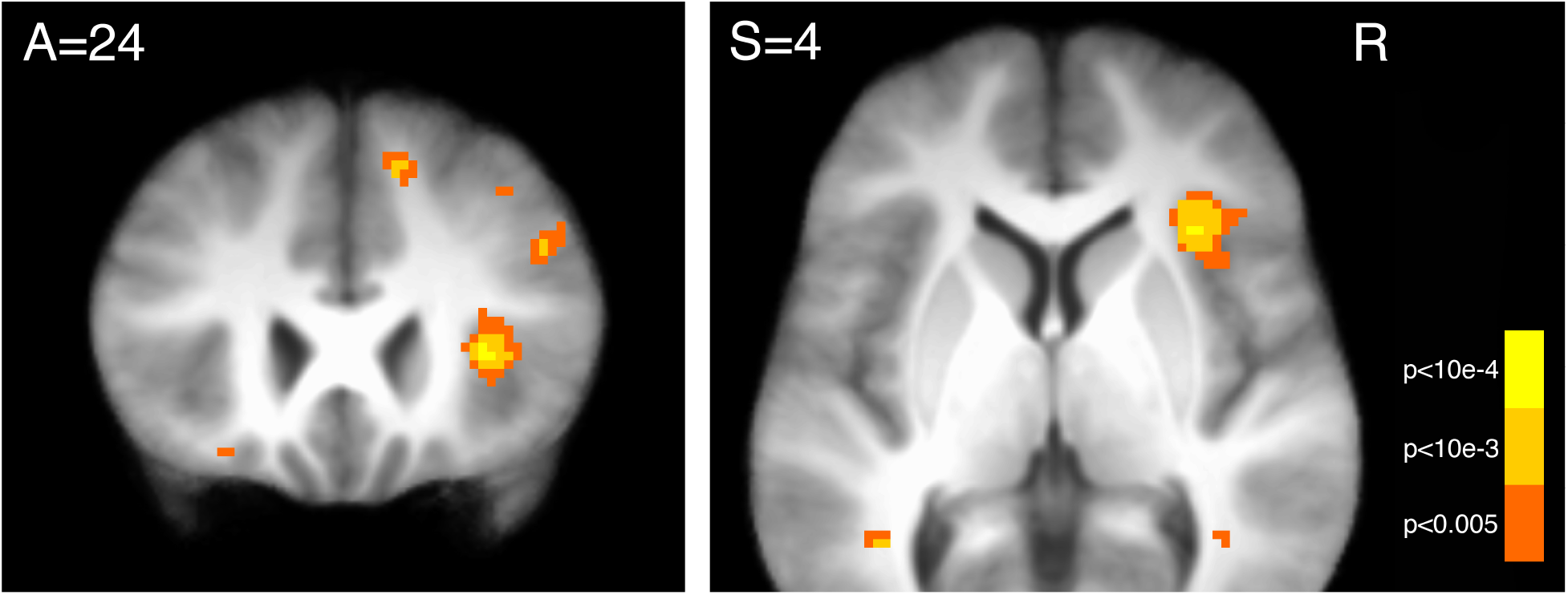
Insula activity during the immediate memory session correlates with memory strength for high-pain associated objects one year later. A contrast of one-year later memory strength for high-pain associated objects vs. memory strength for low-pain associated objects was significantly correlated with activation in the right anterior insula. (Images displayed at p < 0.005, uncorrected for display.)

Next, we examined memory for pain-associated objects using representational similarity analysis (RSA; Kriegeskorte et al., 2008). Previous studies have shown an association between higher within-item similarity and memory (Xue et al., 2010; Ritchey et al., 2013). Building on this within-item similarity memory effect, we predicted that higher across-item distinctness (or dissimilarity) may be related to better memory. In the anterior insula, we indeed found a significant difference in representational similarity, such that patterns for remembered objects were less similar to patterns for other remembered objects than patterns evoked by subsequently forgotten objects (high pain remembered 0.082 ± 0.008; high pain forgotten 0.095 ± 0.009; t(23) = 2.50, p = 0.020). The pattern similarity effect was selective to memory for high pain-associated objects and showed a significantly stronger effect than similarity across low pain objects (high vs. low memory effect comparison, t(23) = 2.08, p = 0.048; low pain memory effect, t(23) = 0.34, p = 0.73). Moreover, the representational similarity difference was not driven by univariate activation, as demonstrated in a regression controlling for trial-by-trial activation in the anterior insula (similarity regression coef. −1.98 ± 0.081; t(22) = −2.45, p = 0.015; bilateral anterior insula coef. 0.18 ± 0.09; t(22) = 1.90, p = 0.058). We found no memory-related differences in the MTL, and when conducting the same analysis using immediate recognition memory strength, we found no difference in the anterior insula (t(_23_) = 0.80, p = 0.44) or MTL.

## Discussion

We found that single episodes incidentally associated with painful experiences were not differentially remembered immediately but showed significantly enhanced memory one year later. We also demonstrate a novel neural correlate predictive of very long-term memory: activity in the anterior insula predicted the strength of memory for pain-associated objects one year later. Further, multivariate patterns of activation in the anterior insula were also related to year-later memory for pain-associated objects.

Our results suggest a mechanism by which memory for neutral information can be enhanced by pairing learning with an arousing experience. Remarkably, the memory enhancement that we observed was for pictures of everyday objects. With only one pairing with an aversive heat pain stimulus, these objects showed better memory that lasted at least one year, even though participants were likely exposed to many of these objects in the real world in the intervening time. Speculatively, the arousal-related memory enhancement we observed may be even more robust for more unique experiences or intentionally studied information.

Neurally, our results suggest a mechanism by which single affective experiences modulate very long-term memory. We found that activity in the anterior insula during the immediate memory test positively correlated with memory strength for pain-associated objects one year later. Paralleling the lack of an immediate behavioral effect of pain on memory, activity in the insula was unrelated to immediate memory strength for high-pain associated objects. However, using univariate and multivariate measures, we found that insula activity was significantly related to memory one year later for high-pain associated objects. During the incidental learning phase, the same anterior insula region was strongly correlated with pain ratings and administered temperature. The anterior insula has been associated with many processes in the fMRI literature, but in the context of pain, it is hypothesized to play a particular role in the emotional and evaluative aspects of pain (Kurth et al., 2010; Wiech et al., 2014). Interestingly, insula activation in post-traumatic stress disorder has also been associated with recollecting traumatic memories (Liberzon and Martis, 2006). The current results suggest that the anterior insula may be related to long-term memory for aversive experiences in healthy human participants. It is possible that the insula activity we observed in the test phase reflects already-engaged memory consolidation processes which lead to very long-term memory benefits for pain-associated objects.

Previous fMRI studies in humans have emphasized the role of MTL-amygdala activity and connectivity in modulating emotional memory over time (Ritchey et al., 2008; Murty et al., 2010; Qin et al., 2012). The amygdala may play an important role in consolidation by triggering the release of neuromodulators such as norepinephrine and corticosteroids (McGaugh, 2013). In our study, heat pain itself elicited activation in the dorsal amygdala / sublenticular extended amygdala. However, we did not find any correlates of immediate or year-later memory in the amygdala, but null effects should be interpreted with caution. It is possible that for aversive somatosensory stimulation, consolidation is related to interaction of the MTL with different regions such as the insula.

As the majority of research on the emotional modulation of memory has utilized emotional pictures, it has remained largely unknown whether and how inherently neutral stimuli may be enhanced by association with an emotional experience (Phelps et al., 1997; Maratos and Rugg, 2001; Smith et al., 2004a; Anderson et al., 2006). When considering inherently emotional stimuli, there does not need to be a memory advantage in order for an agent to act to quickly avoid these stimuli in the future: for example, a large snarling dog remains aversive, and it would be simple to avoid such a threat, even without memory (Phelps et al., 1997; Maratos and Rugg, 2001). Thus, it is important to demonstrate that long-term memory is enhanced for the stimuli that are not so easy to subsequently discriminate and act upon. Increased memory for emotion-associated neutral stimuli would allow for adaptive processing such as increased attentional orienting, which could facilitate more rapid adaptive responding if these stimuli are encountered again.

In conclusion, we demonstrate that a single affective experience increases memory for neutral items one year later. Importantly, our neural results establish a novel connection between brain activity during the initial experimental session and memory one year later, such that increased insula activity predicted later memory strength. While our results were in the negative affective domain, it is possible that a similar memory enhancement would be found for stimuli associated with positive affective experiences. The long-term memory enhancement of neutral elements from emotional experiences may have implications for the understanding and treatment of mood disorders and post-traumatic stress disorder (Hamilton and Gotlib, 2008; Shin and Liberzon, 2010). Further, while our results were for negative arousing experiences, they suggest that positive arousing experiences may also be useful for enhancing learning.

## Author Contributions

G.E. Wimmer and C. Büchel designed the experiment; GEW collected and analyzed data; GEW and CB wrote and revised the manuscript

## Acknowledgments

This work was supported by ERC-2010-AdG_20100407 and DFG SFB TRR 58 and SFB 936. We thank Lea Kampermann for essential assistance with data collection and translation

## Supplementary Information

### Supplementary Results

**Incidental learning phase behavior**. In the incidental learning phase, we administered high and low levels of heat during the presentation of incidental object pictures. Pain ratings given after each trial reliably differentiated high and low heat (high, 7.34 ± 0.06 (mean ± SEM); low, 2.34 ± 0.12; scale range: 1-8). The temperature on high heat trials was on average 49.4 ± 0.3°C and on low heat trials was on average 42.2 ± 0.3°C. Note that temperatures were adjusted on a trial-by-trial basis to maintain a difference in pain ratings. On the 6 trials where visual cue-pain association knowledge was assessed, the heat level associated with cues was correctly identified on 89.9 ± 4.8% of probes.

A pain-predictive cue preceded the onset of heat and the incidental picture; on some trials the heat expectation created by the predictive cue was violated. Expectation violation tended to increase pain ratings for low heat trials where the expectation was for high heat (high expectation and low heat vs. validly cued low heat: t(_23_) = 1.90, p = 0.07; low expectation and high heat vs. validly cued high heat: t(_23_) = 0.67, p = 0.51). These results indicate that the high and low levels of pain were clearly discriminable and that expectation had little effect on pain experience.

**Memory effect of visual cue expectation violation**. On the trials where cue-induced heat expectations were violated, unexpected high pain resulted in numerically lower immediate memory (regression analysis, t(_21_) = −1.22, p = 0.22; unexpected low vs. expected low, t(_21_) = 1.28, p = 0.20). At the year-later memory test, similar to the numerical effects in the immediate test, we found overall decreased memory for objects associated with unexpected high pain vs. expected high pain (regression analysis, t(_22_) = −2.50, p = 0.013; unexpected low vs. expected low, t(_22_) = −1.18, p = 0.24).

**fMRI results**. In the fMRI analysis, prior to memory analyses we verified the main effect of high vs. low pain. During the incidental learning phase, trial-by-trial pain ratings positively correlated with activation in a wide system of regions previously implicated in pain processing (Apkarian et al., 2005) including the anterior and posterior insula, cingulate, thalamus, and secondary somatosensory cortex (all p < 0.05 whole-brain FWE corrected; Fig. S2 and Table S1). In a region-of-interest analysis in the MTL (including the hippocampus, parahippocampal cortex, and the amygdala), we found bilateral activation in a region consistent with the dorsal amygdala / sublenticular extendend amygdala (Table S1).

In the incidental learning phase, at the onset of the object stimuli and heat, overall immediate test memory strength was positively correlated with activity at an uncorrected level in the left hippocampus (−24, −10, −22; Z = 2.88, p = 0.002, unc.) as well as in the right posterior occipital cortex (34, −78, 8; Z = 3.31, p < 0.001, unc.), a region also activated by presentation of object stimuli. These correlates of immediate memory at object onset in the incidental learning phase were not present when we looked at later activity during the peak pain period (5 s later; see Supplemental Experimental Procedures).

**Fig. S1.**
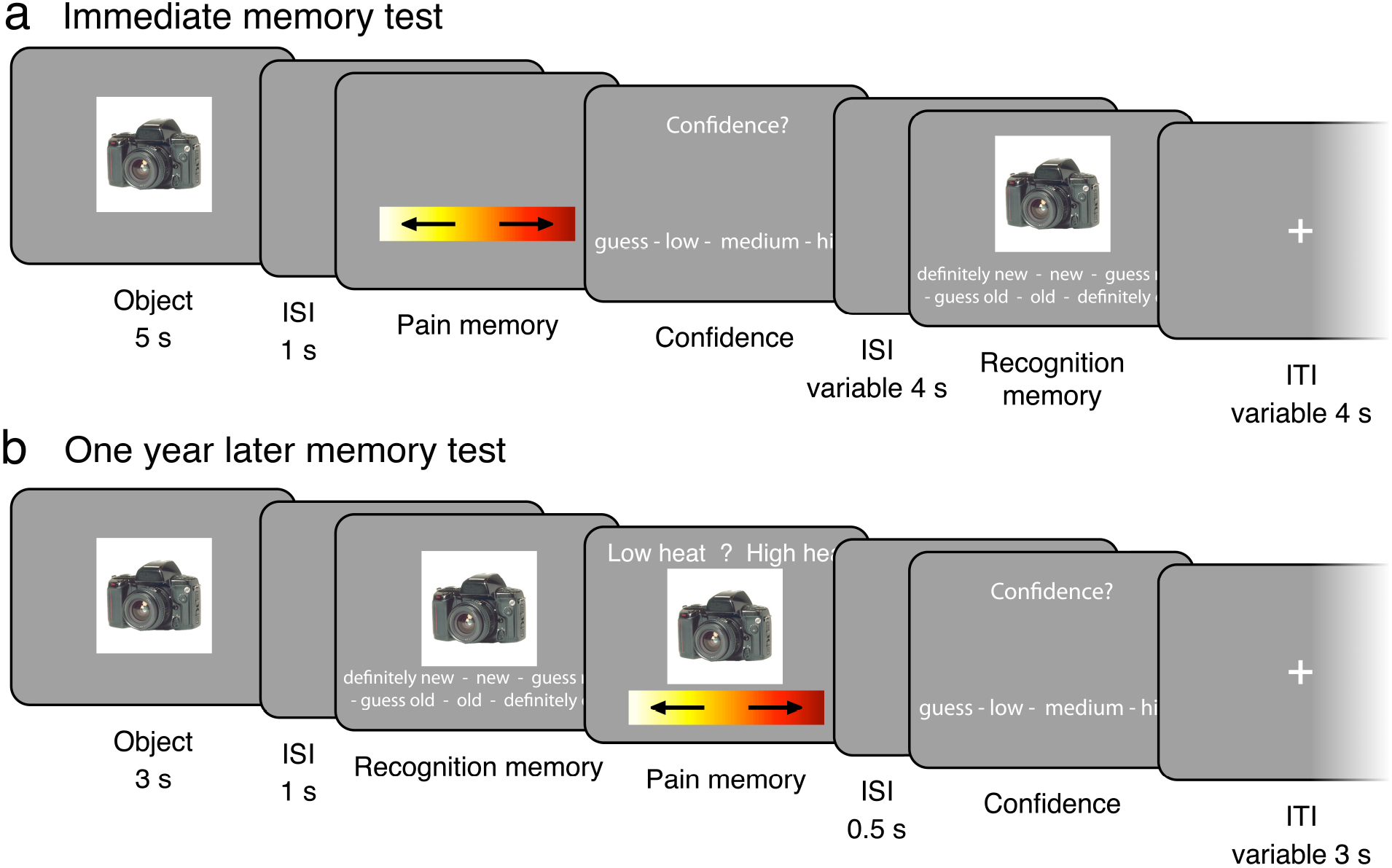
Immediate and one year later surprise memory tests. (a) Immediate memory test phase. After viewing an incidental object from the learning phase, participants responded with whether the object was associated with high or low pain and then rated their confidence in this response. Participants then rated their recognition strength on a 6-point new-to-old scale. (b) One year later memory test phase. After viewing an incidental object from the learning phase, participants rated their recognition strength on a 6-point new-to-old scale. If the object was rated “old”, participants then responded to a binary pain memory question about whether the object had been associated with high or low pain in the incidental learning session one year prior, and then rated their confidence.

**Fig. S2.**
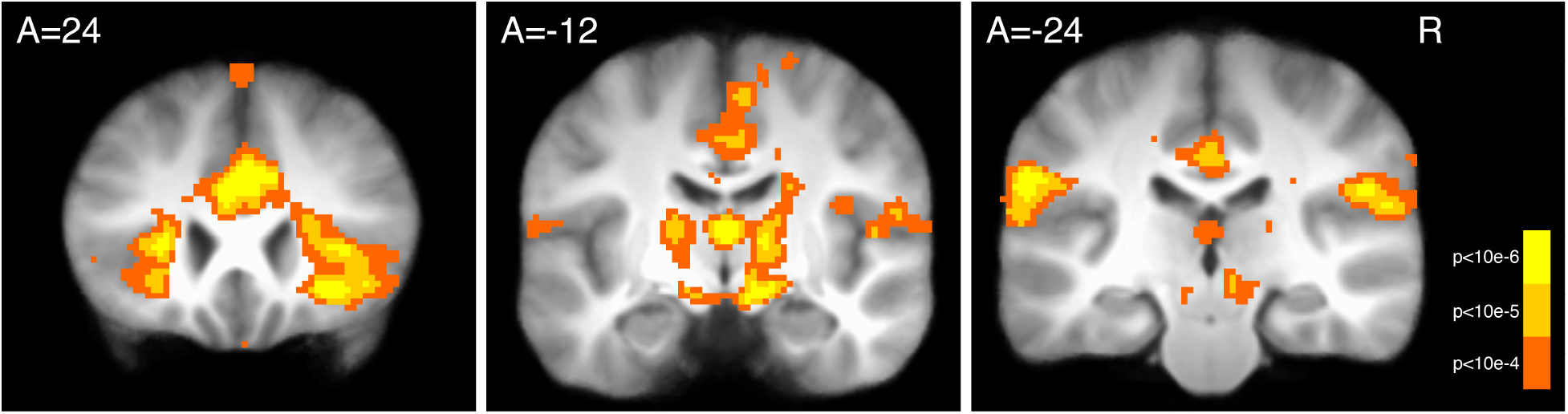
Pain-correlated responses during the incidental learning phase. Brain activation was positively correlated with trial-by-trial pain ratings in the anterior insula, cingulate (left panel), thalamus, midbrain (middle panel), and secondary somatosensory cortex (right panel). See also Table S1. (Images thresholded at p < .0001 uncorrected for display; A = Anterior, R = Right.)

**Table S1.**
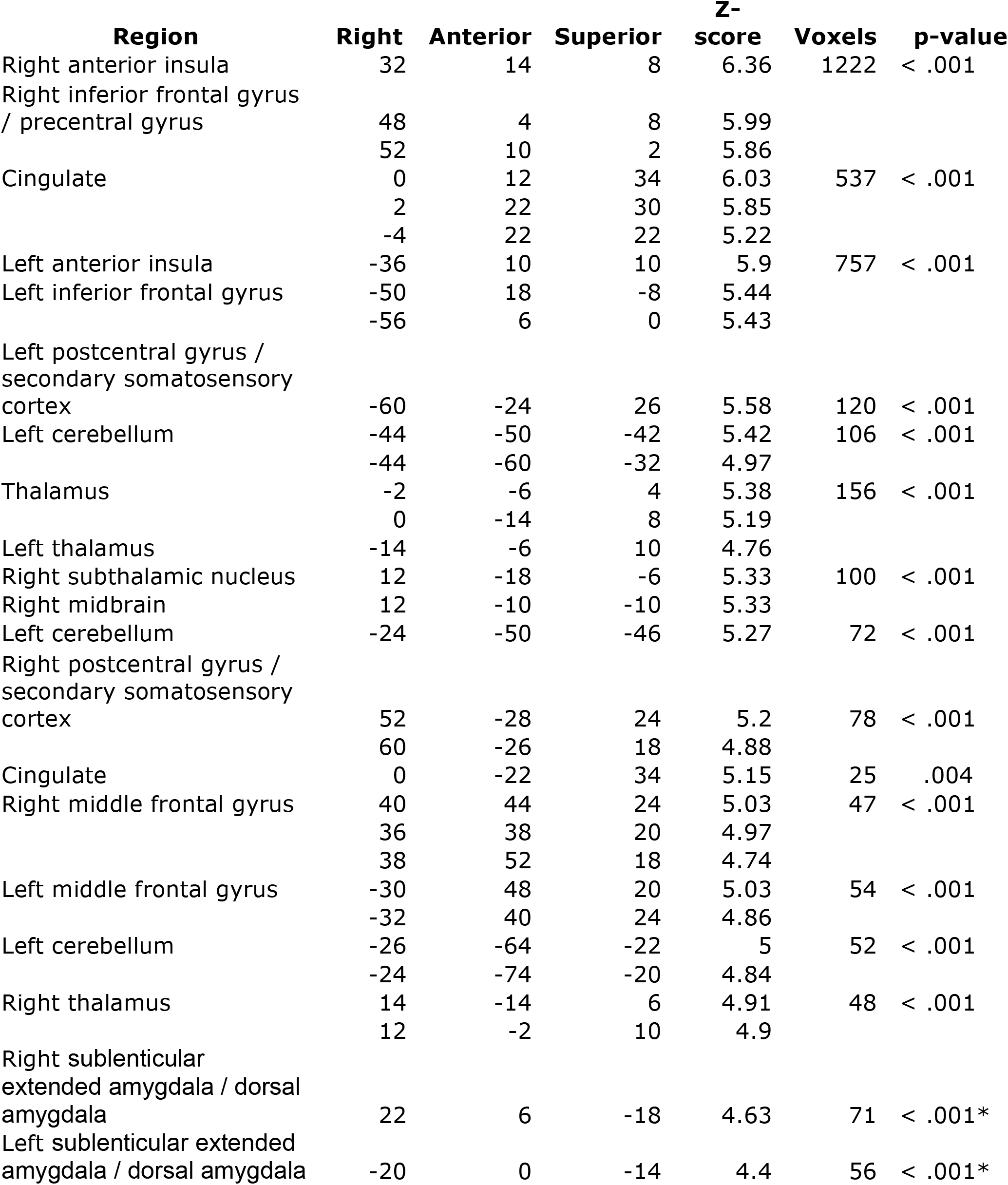
Neural correlates of pain ratings during pain administration in the incidental learning phase. All p-values are whole-brain FWE-corrected, except where * indicates SVC p-values.

| Region | Right | Anterior | Superior | Z-score | Voxels | p-value |
| --- | --- | --- | --- | --- | --- | --- |
| Right anterior insula | 32 | 14 | 8 | 6.36 | 1222 | < .001 |
| Right inferior frontal gyrus / precentral gyrus | 48 | 4 | 8 | 5.99 |  |  |
|  | 52 | 10 | 2 | 5.86 |  |  |
| Cingulate | 0 | 12 | 34 | 6.03 | 537 | < .001 |
|  | 2 | 22 | 30 | 5.85 |  |  |
|  | -4 | 22 | 22 | 5.22 |  |  |
| Left anterior insula | -36 | 10 | 10 | 5.9 | 757 | < .001 |
| Left inferior frontal gyrus | -50 | 18 | -8 | 5.44 |  |  |
|  | -56 | 6 | 0 | 5.43 |  |  |
| Left postcentral gyrus / secondary somatosensory cortex | -60 | -24 | 26 | 5.58 | 120 | < .001 |
| Left cerebellum | -44 | -50 | -42 | 5.42 | 106 | < .001 |
|  | -44 | -60 | -32 | 4.97 |  |  |
| Thalamus | -2 | -6 | 4 | 5.38 | 156 | < .001 |
|  | 0 | -14 | 8 | 5.19 |  |  |
| Left thalamus | -14 | -6 | 10 | 4.76 |  |  |
| Right subthalamic nucleus | 12 | -18 | -6 | 5.33 | 100 | < .001 |
| Right midbrain | 12 | -10 | -10 | 5.33 |  |  |
| Left cerebellum | -24 | -50 | -46 | 5.27 | 72 | < .001 |
| Right postcentral gyrus / secondary somatosensory cortex | 52 | -28 | 24 | 5.2 | 78 | < .001 |
|  | 60 | -26 | 18 | 4.88 |  |  |
| Cingulate | 0 | -22 | 34 | 5.15 | 25 | .004 |
| Right middle frontal gyrus | 40 | 44 | 24 | 5.03 | 47 | < .001 |
|  | 36 | 38 | 20 | 4.97 |  |  |
|  | 38 | 52 | 18 | 4.74 |  |  |
| Left middle frontal gyrus | -30 | 48 | 20 | 5.03 | 54 | < .001 |
|  | -32 | 40 | 24 | 4.86 |  |  |
| Left cerebellum | -26 | -64 | -22 | 5 | 52 | < .001 |
|  | -24 | -74 | -20 | 4.84 |  |  |
| Right thalamus | 14 | -14 | 6 | 4.91 | 48 | < .001 |
|  | 12 | -2 | 10 | 4.9 |  |  |
| Right sublenticular extended amygdala / dorsal amygdala | 22 | 6 | -18 | 4.63 | 71 | < .001* |
| Left sublenticular extended amygdala / dorsal amygdala | -20 | 0 | -14 | 4.4 | 56 | < .001* |

